# Ovule development in Cycads: observation on anatomy and nucellus morphology in *Zamia* and *Cycas*

**DOI:** 10.1101/735837

**Authors:** Xin Zhang

## Abstract

In the ovule evolution, the integument is the most attention point in discussion as a morphologic character of the seed plants. There are several theories and hypotheses about the origin of the integument were presented in the history. However, the development and function of the ovule envelopes are not so clear until now. The development of thehe basal gymnosperms Cycas and Zamia were to investigated, especially of the integument to complement the existing knowledge in seed plants. The development of ovules of seed plant is documented with morphological and anatomical using LM and SEM.

The nucellar beak found in Zamia is a structure that has not been recorded previously. It protrudes from the micropyle at pollination and may be the primary acceptor for pollen. There are striking similarities to the lagenostom or salpinx in Lyginopteridatae. There may be an evolutionary way to interpret the pollination drop existing in the Lyginopteridatae. Probably the nucellar beak of Cycads, even Ginkgoales have the same function with the lagenostom or salpinx of the Lyginopteridatea. Unfortunately, pollen and transport inside the pollination chambers have not been observed. Further analysis of this unusual structure seems to be very important.

## Introduction

Cycads are well studied in respect of morphology and anatomy. However, in conifers freely exposed pollination droplets are well documented. While similar records are lacking for Cycads. There must be a liquid phase involved as spermatozoids would not swim otherwise. On the other hand, artificial pollination under greenhouse conditions are performed in *Zamia* by rinsing water with dispersed pollen through the strobilus. Freely exposed pollination droplets would be washed away applying this method. The method is working successfully, but this is in conflict without present knowledge of the pollination procession of Cycads. A more detailed analysis is thus essential.

“Interest in the morphology of the ovules of living Cycads has been much stimulated by the recent appearance of several papers dealing with the structure of Cycad-like seeds of Carboniferous age. The structures found in these fossils are not only of great interest in themselves, but also seem to throw light on some of the difficult points in the morphology of recent seeds, and it became clear that a detailed examination of the living forms might be undertaken with advantage” (Stopes, 1905).

The ovule of seed plants is the female structure with at least one or two envelop structures covering the central part called nucellus or megasporangium. The only point of agreement in the different theories on the origin of ovule is that the entire nucellus is a megasporangium that retains a single megaspore and the endospermic female gametophyte (HerrJr, 1995).

During the latter part of the nineteenth century several conflicting theories based on anatomical, developmental and teratological evidence from extant plants were put forward, but no definite conclusion was reached. Worsdell (1904) discussed the three principal theories for ovule origin advanced during the nineteenth century namely the axial theory, the foliolar theory and the sui generis theory. The axial theory regards the nucellus as a shoot axis bearing foliolar appendages fused to form the integument. This is supported by the morphology of gymnosperms, especially the strobilus structure of conifers. The foliolar theory held the ovule to be morphologically a phyllome; it relates the ovule to a three-lobe leaflet of the female sporophyll (or carpel in the angiosperms); the integument comes from the lateral ones; the nucellus is an emergence borne on the cup-shaped terminal lobe. The foliolar theory has various forms. It is supported by the angiosperm ovule particularly (Eames, 1961). The sui generis theory is that the ovule is a new structure which is not homologous to either caulome or phyllome. These three theories generally lost influence. However, one can also find their shadow in some of the botanical text books today.

In the 20th century the consideration of the ovule origin has focused largely on Pteridospermophyta, which are generally accepted to include the oldest seed plants. Benson (1904) noticed that the synangium of *Telangium scottii* Benson and the ovule of *Lagenostoma* are very similar to each other. She hypothesized that the integument evolved from a sterile outer ring of sporangia in a radial synangium. This is the first theory of the pteridosperm ovule based on fossil evidence. According to her theory, the number of spores in the central sporangium was reduced to one which finally produced the female gametophyte.

Oliver and Scott (1904) hypothesized the evolution of the integument from a cup-like indusium in their monograph on *Lagenostoma lomaxii*. Regarding the origin of the ovule they stated that “we have in *Lagenostoma* a megasporangium which has been enclosed by two successive, concentric, indusium-like structures of which the inner has become an integral part of a new organ, the seed; the outer is probably of later origin. It is quite possible that the two enclosures have originated very similarly, i.e. as peltate, lobed structures, and that the present integument was once a comparatively unspecialized, cupule-like indusium”. In the monograph on *Lagenostoma* they state, “a comparison of the seeds of Cycads with *Lagenostoma* is inevitable”. Stopes (1904, 1905) and Matte (1904) supported this idea with their work on the integument structure of the Cycads. They showed independently that the inner vascular system of the cycad ovule supplies the inner part of the integument but not the nucellus as previously thought, and the integument itself has a double structure in respect to vascularization. Stopes considered that the integumentary vascular system of *Lagenostoma* is equivalent to the inner system of Cycads and that the cupular system is equivalent to the outer Cycads system. The integument of *Lagenostoma* is thus regarded as homologous with the inner layer of the Cycads integument and the cupule is homologous with the outer part. As the three papers (Oliver and Scott 1904; Stopes 1904; Matte 1904) dealing with Cycads and *Lagenostoma* appeared simultaneously, the comparison does not cover all details. Stopes (1905) stated “I will not attempt to do this now in detail, but there are one or two points about which I should like to add something to my published view”. Work on the anatomy and morphology of living Cycads revealed that their integumentary structure is more complex than was generally supposed (Stopes, 1904). In fact, that this comparison of Cycads and *Lagenostoma* was not been done until now.

The investigations concerning the ovule started from the conditions of the angiosperms and to state the nature of nucellus and integument, five different approaches were used, ontogeny, anatomy, topographical-morphology, phylogeny and teratology (de Haan, 1920). These units are now referred to as telomes base on the idea of de Haan (Smith, 1964; HerrJr, 1995). Zimmermann (1938, 1952) developed this concept of telomes which conceived a dichotomously branched axial system bearing terminal sporangia as the evolutionary starting point. The ultimate axes are termed telomes, either sterile or fertile. Parts between to branching points are called mestomes. Through time there is a gradual reduction of some of the axes so that a single sporangium becomes surrounded by an aggregation of sterile processes. Several Devonian-Mississippian seeds exhibit some of the structural modifications suggested by the telome theory, as the earliest integuments are formed of unfused axes surrounding the nucellus. This theory has been revised by several investigators (Walton, 1953; Andrews, 1961; Smith, 1964; Long, 1966; Gensel and Andrews, 1984; Taylor and Taylor, 1993; HerrJr, 1995; Taylor, Taylor and Krings, 2009), and the concept is regarded as the most plausible to date (HerrJr, 1995). Smith (1964) considered it to be the only theory to account for the structural variations in pteridosperm ovules. However, there are two things that should be noticed. The first step of this concept is hypothetic; second, what is the structure of the telome and how this structure was evolved to fit the function of pollination and fertilization is not clear. The telome concept is widely considered the most plausible one. Andrews (1961) reviewed the theories on the evolution of the ovule and considered that except the telomic concept there is another principal theory, the nucellus modification concept. Andrews (1961) starts the evolutionary series from the megasporangium of the Lower Carboniferous fern *Stauropteris burntislandica* Bertrand which has only two functional megaspores. The tapering apex of the sporangium often has a small pore and the sterile lower half is traversed by a central vascular strand.

Walton (1953) hypothesized that a portion of a frond or a leaf became enrolled, based on the presence of vascular tissue in the peripheral nucellus of Trigonocarpels, which he considered as an integument fused to a strongly reduced nucellus. He regarded the previously designated integument as equivalent to the cupules of *Lagenostoma*.

Camp and Hubbard (1963) proposed the double integument theory by which Paleozoic ovules were regarded as having two adnate integuments derived from sterile dichotomous lateral branches subtending a terminally borne megasporangium. The inner set of branches fused with the nucellus so that the ovules appear to have only one integument. From a comparison of pteridosperm and *Cycas* ovules with those of angiosperms, they suggested a greater homology among ovules of different groups that all ovules have two integuments whether separate or fused. Meeuse and Bouman (1974) also set a similar hypothesis.

The pollination drop was present even in the Paleozoic times (Singh, 1978). The most obvious adaptive significance of an integument appears to be an increased protection of the megagametophyte and embryo, but the free integumentary lobes of the earliest seeds may also have served to increase pollen capture (Taylor, Taylor and Krings, 2009).

Normally, fertilization is the key point of the development from the ovule to seed. Prior to fertilization, the ovule develops. The covering sheath is called integument. After fertilization, the ovule matures to form seed. The integument becomes the seed coat or testa. However, compared to seed development fertilization is just a very short time and in the development from the ovule to the seed, there is no obvious interruption. So here the development of ovule is the development from the primordium to the immature seed. The covering structure is the integument even in young stages of a seed after the fertilization.

## Material and methods

Different developmental stages of inflorescences were collected from the Botanical Garden of the Ruhr University Bochum in 2010 and 2012, fixed in FAA (Formalin: Acetic acid: Alcohol 70% = 5:5:90) and kept in the fixative under moderate vacuum for at least 30 min to optimize infiltration of the fixative. After 2 days the FAA was replaced by 70% ethanol for further storage.

Dehydration of the dissected flowers was performed using FDA (Formaldehyde Dimethyl Acetal) for 24 hours. For critical point drying a BALZERS CPD 030 was used. The samples were mounted on aluminum stubs and sputter coated with gold for 200 seconds at 42-43 mA.

SEM studies were performed with a ZEISS DSM 950 SEM. For documentation, the digital recording system "Digital Image Processing System 2.2" was used at a resolution of being 2000 × 2000 pixels, to avoid loss of detail from the original image content.

Anatomical studies were done using classical paraffin serial sectioning with a Leica RM 2065 rotary microtome, sections from 5-8 μm were stained using astramin/astra-blue staining, Photographs were made with a Zeiss Axioplan supplied with a color view II camera (Olympus). For multiple image alignment the software Cell F (Olympus) was used. All figures are organized for publication using Photoshop 7.0.

## Results

### The morphology of Zamiaceae

*Zamia amblyphyllidia* D. W. STEV (Fig. 1) is a representative of the Cycadales, an evolutionarily ancient gymnosperms order. They are generally known as cycads having the original features of fossil gymnosperms, like all other representatives of the order. First, the megasporophyll is a leaf like structure (Fig. 1). Second the sperm is a spermatozoid, what in addition to Cycadales is only found in *Ginkgo biloba* L. Third, like many Cycadales, they resemble tree ferns strongly in appearance because of their leaf crown, with large fronds (Fig. 1, Fig. 9). *Z. amblyphyllidia* is pachycaul due to the branching directly above the ground. Distally at each end of the branched trunk the leaves form a crown with a maximum of 15 leaves that can reach a length of 150 cm and up to 40 pairs of uniformly distributed leaflets. The individual leaflets are narrow-lanceolate and slightly serrate in the distal quarter. Samplestaken from *Zamia amblyphyllidia* come from the Botanical Garden of the University of Bochum. The plants are kept in pots in a temperate greenhouse (Fig. 1). There are several female and male individuals of *Z. amblyphyllidia*. In the vegetative stage both sexes cannot be distinguished, unlike *Cycas* the strobilus is grouped in 2-5 and is not proliferating. The slender strobilus of male plants (Fig. 2) is cylindrical in shape and can grow up to 8 inches long. For pollination the red-brown sporophylls diverge. After pollen shed the male strobilus dies.

**Fig. 1:**
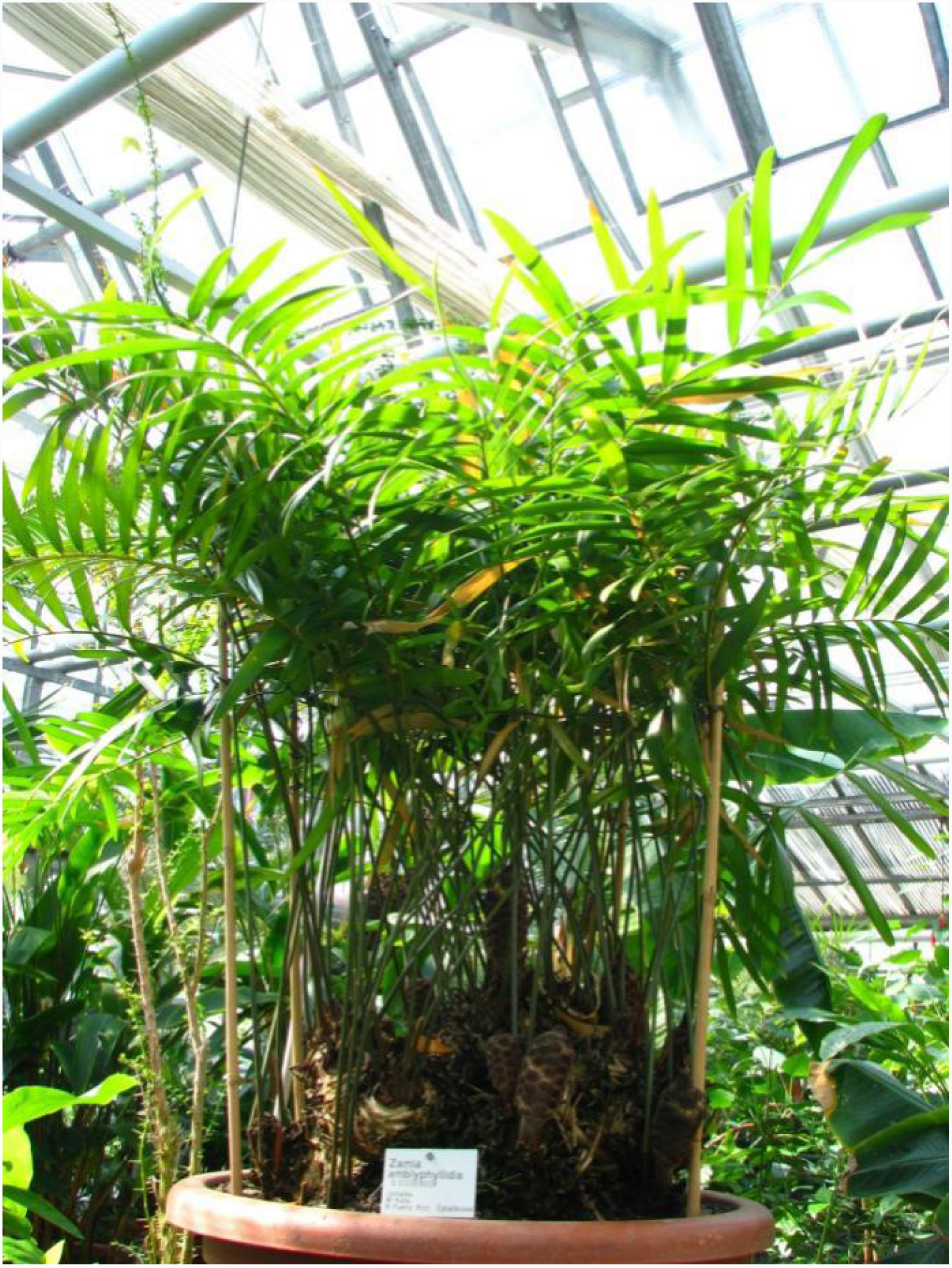
The plant of *Zamia amblyphyllidia* D. W. STEV.

The much more compact, cylindrical female strobilus (Fig. 3) is about 15 cm long and reaches a diameter of 4-6 cm. The megasporophylls form a compact strobilus and have a well-defined stalk which expands into a peltate head. The cone has a slightly ridged apex (Fig. 3 arrows). At pollination the apex is dark brown (Fig. 3 green arrow) and later rather grayish (Fig. 3 red arrow). The individual sporophyll separates for pollination just below the cone apex and near the cone basis. After the pollination, the female strobilus closes again and remains extremely compact during the entire maturation. The cone scales are forced apart only by the increasing seeds.

**Fig. 3:**
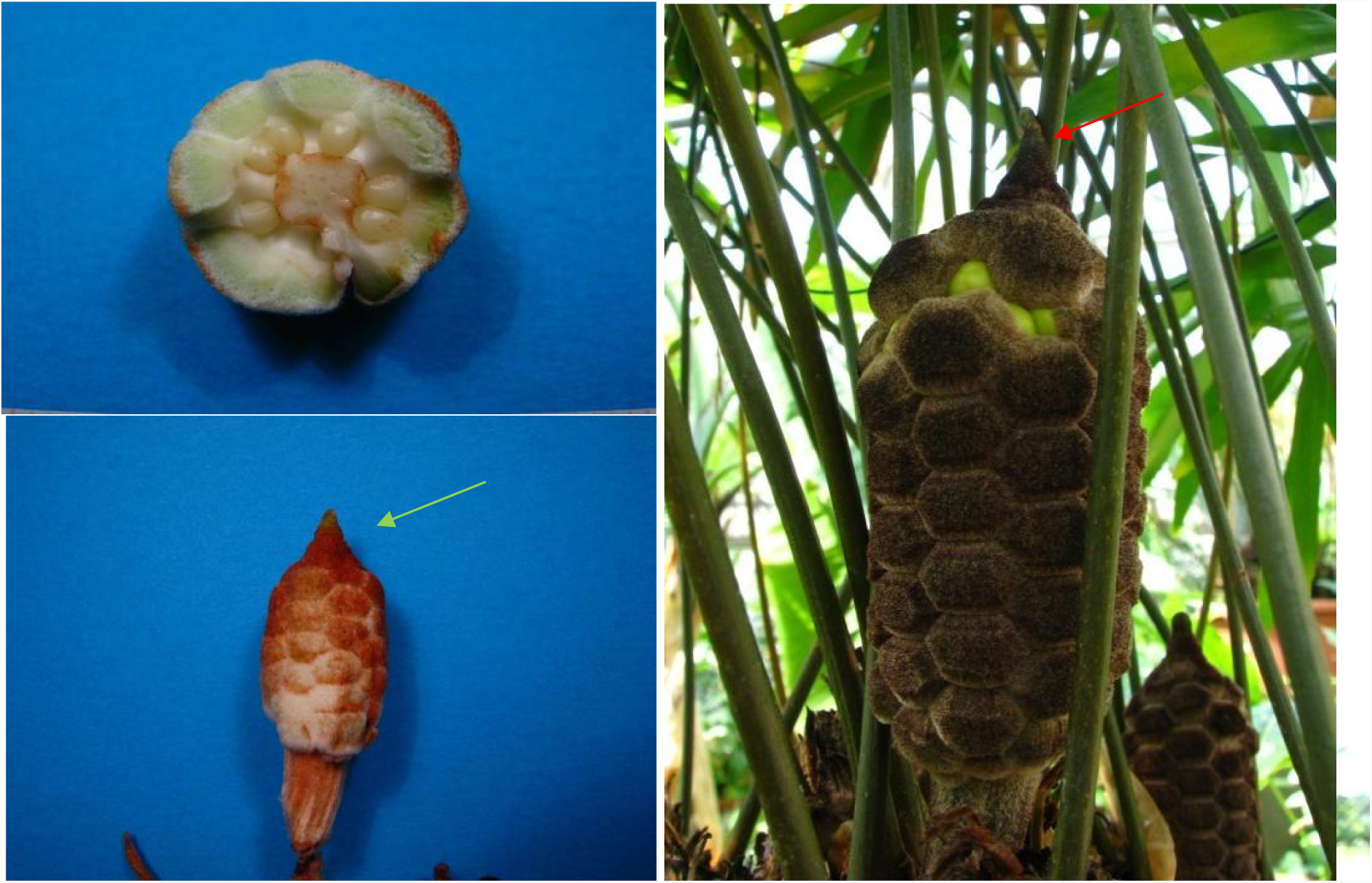
The female strobilus of *Zamia amblyphyllidia* D. W. STEVyoungerstage and mature stage.

As in the Botanical Garden Bochum several female specimens are available, cones were collected depending on arrange in size from different plants similar in size and age. To obtain a complete developmental sequence fixed material was used collected from 2009 to 2012.

Normally two ovules are borne on each peltate head which is elongated transversally, and an ovule is borne on either side in the transversal plane (Fig. 4). Occasionally a third ovule may occur. The two ovules are situated one on either side of the short stalk, and their micropyle points radially towards the axis of the strobilus. The ovules are clearly not marginal, but each is situated slightly within the margin of a small area apparently corresponding to one of the areas of the lower surface of a female sporophyll.

**Fig. 4:**
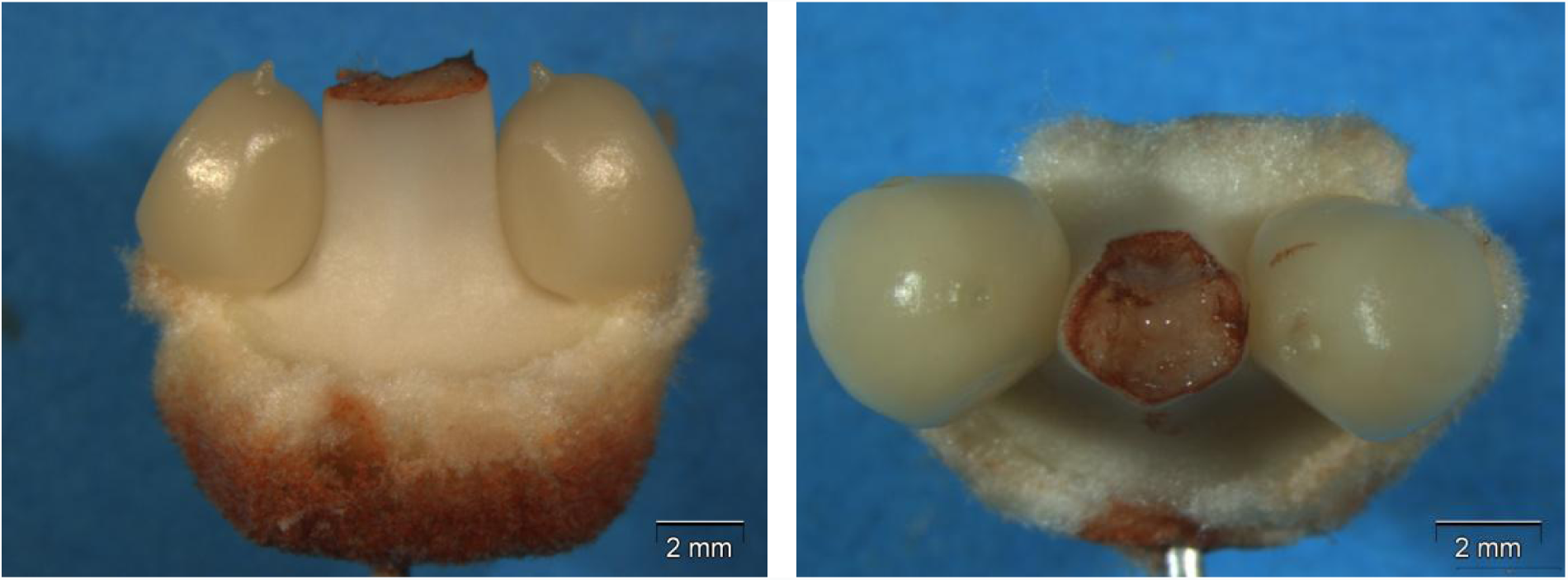
The microsporophyll of *Zamia amblyphyllidia* D. W. STEV.

It is impossible to obtain strobilus which is young enough to show the first stages of the development of the ovule without destroying the plant.

### The ovule development and seed anatomy of *Zamia*

The longitudinal growth of the ovule takes place predominantly in the distal region of the ovule, where the integument is free. From this growth results that the nucellus is fused only to about half its length to the integument before the development of mega prothallium begins. Reinforced by a previous completely extension of the integument growth the integument now encloses the nucellus (Fig. 5A, B). There is a protruding beak on top of the nucellus pointing to the micropyle.It is callednucellar beak. This nucellar beak grows into the micropylar channel and fills it step by step completely (Fig. 5, A, C, E). In parallel the micropylar channel gets closer and closer and the opening finally appears irregularly lobed in top view of the ovule. At this stage, the further differentiation within the nucellus goes on with a size increase of the integument macrospore (Fig. 5E, F).

**Fig. 5:**
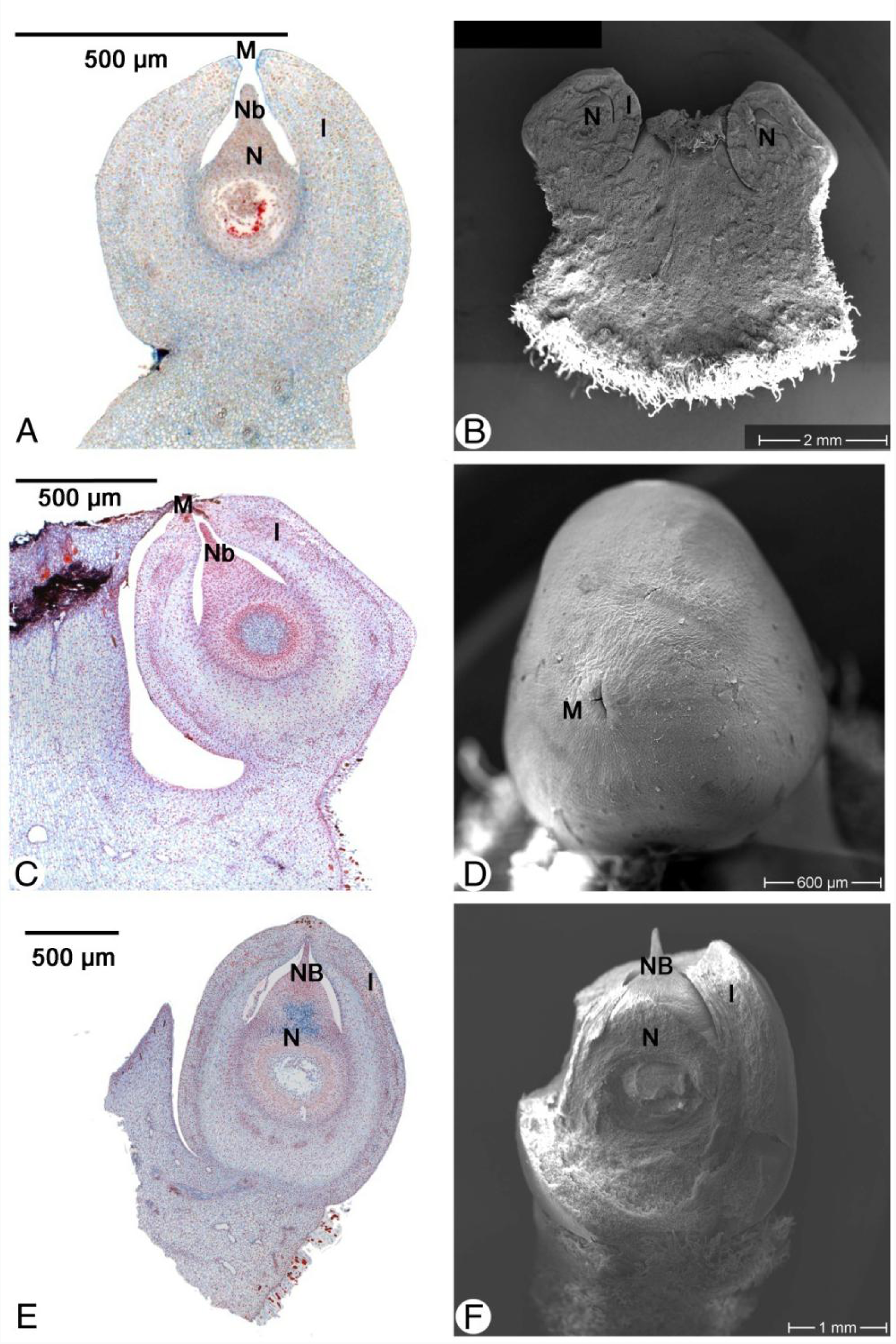
Ovule development I of *Zamia amblyphyllidia* D. W. STEV. A, C and E – The longitudinal sections of the ovule from young to older stage; B, D and F – the same stage of A, C and E with the SEM results; B – the longitudinal section of one megasporophylls with two young ovules; F – the integument is taken partly and the nucellus and nucellus beak can be saw. A, I = Integument, M = Micropyle, N = Nucellus, NB = Nucellar beak.

The integument forms a micropylar tip (Fig. 6A, B and C). The Nucellar beak with longitudinal cells reaches from the tip until the macro prothallium (Fig. 6D, F). The protruding nucellar beak first reaches the middle of the micropylar channel and fills it completely without leaving any space between nucellus and micropylar tissue (Fig. 6E). By further growth the nucellar beak it extends to the distal end of the micropylar channel and can be seen from outside in top view of the ovule (Fig. 7B).

**Fig. 6:**
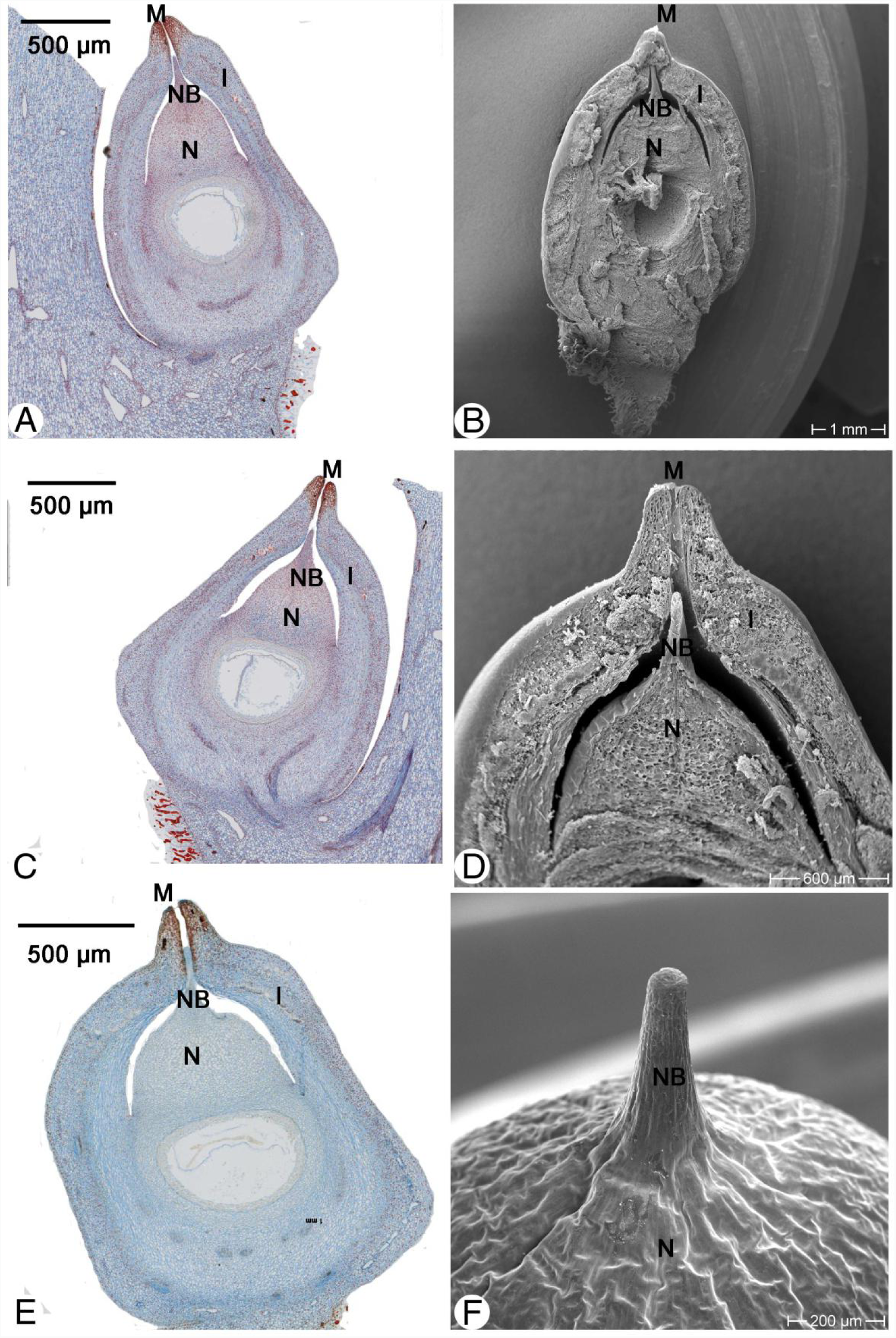
Ovule development IIof *Zamia amblyphyllidia* D. W. STEV. A, C and E – The longitudinal sections of the ovule from young to older stage before the pollination; B, D and F – the same stage of A, C and E with the SEM results; B and D – the longitudinal section to show the position of the nucellus,nucellar beak and the integument; F – the integument is taken to show the morphology of the nucellus beak only. I = Integument, M = Micropyle, N = Nucellus, NB = Nucellar beak.

**Fig. 7:**
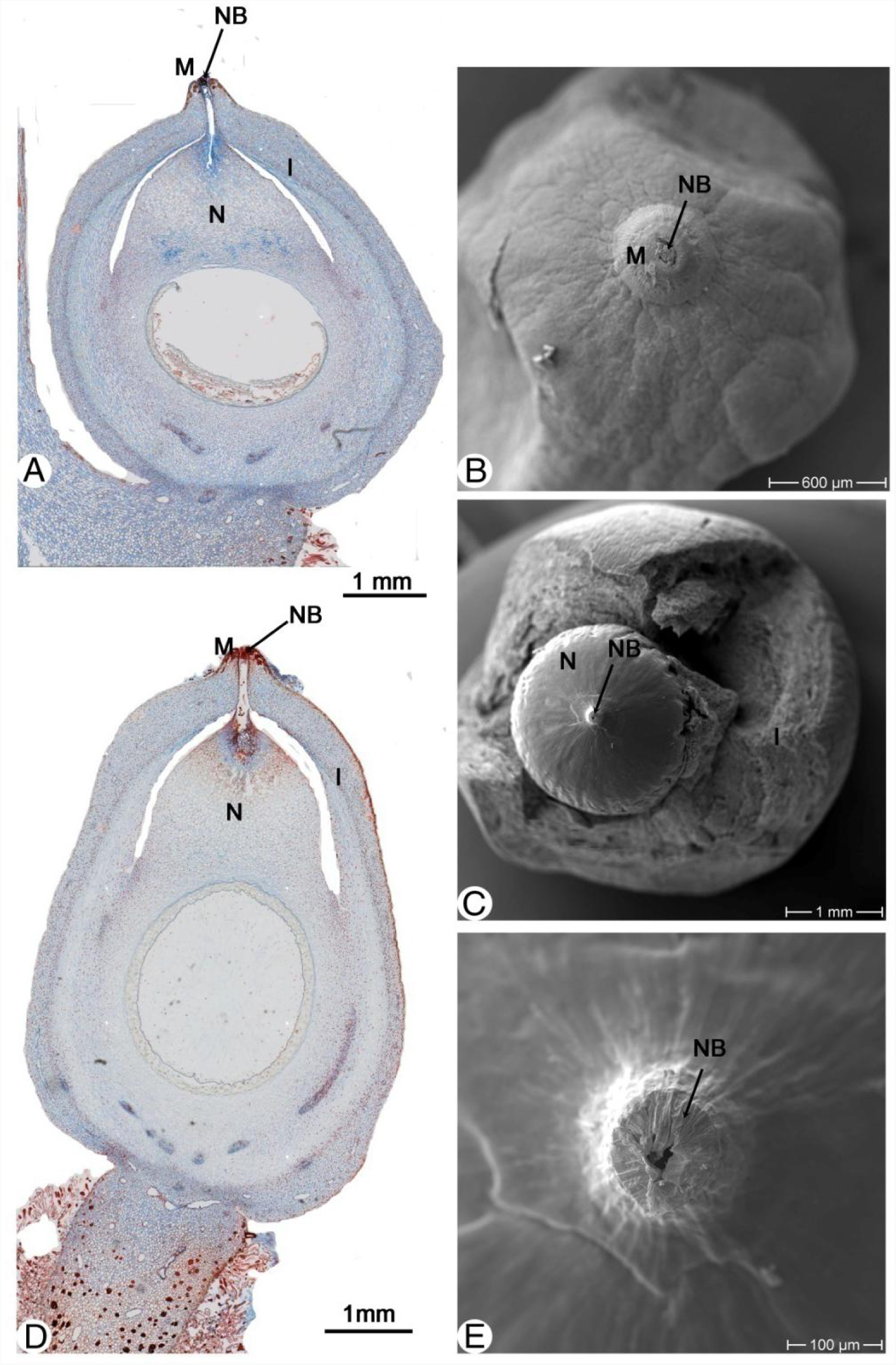
Ovule development IIIof *Zamia amblyphyllidia* D. W. STEV. A, the longitudinal section of the ovule with the nucellar beak reach to the micropyle; B, overlook of the ovule from the micropyle to show the nucellar beak reach to the micropyle, C, the integument is taken to show the nucellus and nucellar beak with a hole on the top of nucellar beak; E, the magnification of the nucellar beak in C with a hole on the top of cellsdeliquesce; D, the longitudinal section of the later stage of A with some pollen in the micropyle canal. I = Integument, M = Micropyle, N = Nucellus, NB = Nucellar beak.

At about this stage the lysis of the tissue of the nucellar beak begins starting from its distal end and proceeding gradually down to the mega prothallium. At early stages the nucellar beak seems to be still closed (Fig. 7A). An exposed pollination drop has never been observed despite seeking carefully for such a stage. Only when the lysis has reached the main body of the nucellus, a widening to the pollination chamber can be seen (Fig. 7D). At this stage the peripheral wall of the nucellar beak is still present forming a kind of chimney that can be detected as a dark blue line in longitudinal sections. It is unclear whether the beak tip is still closed or already open at this stage. In longitudinal sections the distal end of the beak appears blue due to astra blue staining and no clear cell margins are visible. Slightly later the distal end shows a clear opening in SEM samples (Fig. 7C, E).

After pollination, the strobilus is closed again. The pollen inside the nucellus forms the pollen tube and pass the tissues of the chamber between the nucellar beak and macro prothallium until mid-February in the nucellus (Fig. 8A). At this moment, there are several micro prothallia developed in the pollen chamber in different stages (Fig. 7A red arrows). There are two archegonia within the section, each forming neck cells (Fig. 8B). We found a special structure on the top of one archegonium (Fig. 8C, D).

**Fig. 8:**
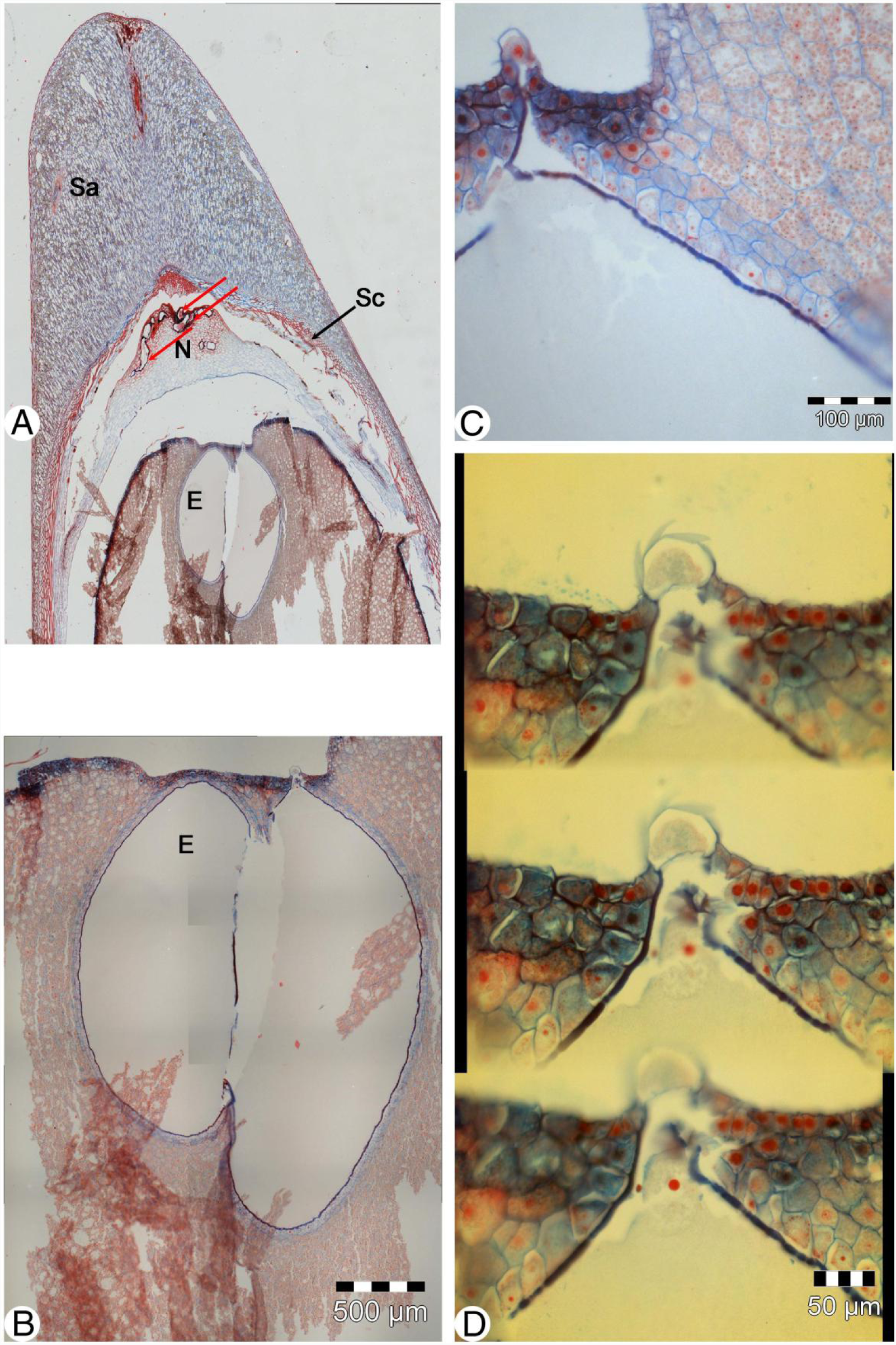
Seedanatomy of *Zamia amblyphyllidia* D. W. STEV.A – the upper part of a mature; B – the magnification of the egg part of A; C and D – to show the detail structure of the archegonium. Red arrows show micro prothallium with in nucellus. E = Egg, N = Nucellus, Sa= Sarcotesta, Sc = Sclerotesta.

### The ovule morphology and anatomy of *Cycas revoluta*

Cycadaceae, described by Carolus Linnaeus in 1753, comprise the only genus *Cycas* which probably consists of about forty species distributed in South-East Asia, southern China, Malaysia, tropical Australia and various islands of the western Pacific, with a disjunction species from Africa and Madagascar (Jones, 1993). Normally, *Cycas revoluta* forms an unbranched monopodial trunk which reaches up to 2.5 m and 30 cm in diameter. The trunk may produce occasionally offsets produced on the trunk or from the base Those offsets do not develop to mature plants. Young leaves are bright green, bearing grey hairs (Fig. 9A and 9B). The growing point of the female plant grows through the developing crown of megasporophylls (Fig. 9C). New leaves are erect with circinate leaflets, emerging with hairs setting lost with age (Fig. 9B). The megasporophylls form a loose and open cone on the top of the trunk (Fig. 9C). It is brown and hairy. The megasporophyll (Fig. 10) of *Cycas* is a distinct foliar organ, divided into lobes, bearing the ovules on its margins. There are more than two ovules on each megasporophyll. The ovule (Fig. 10) is bilateral symmetrical with a length of about 1.5 cm and a diameter of 0.8 cm at anthesis. It is covered near the micropyle with hairs, whereas young ovules are wholly clothed by a dense indumentum except for the micropyle. The ripe ovules are elliptical and can reach a length of about 3 cm and a greater diameter of 2.7 cm. The epidermal cells of the ripe ovules are less radically extended than in other Cycads and develop a strong orange-colored epidermis. The integument clearly shows three different layers (Fig. 11), an outer fleshy, a middle layer of highly thickened, and an inner fleshy layer.

**Fig. 9A:**
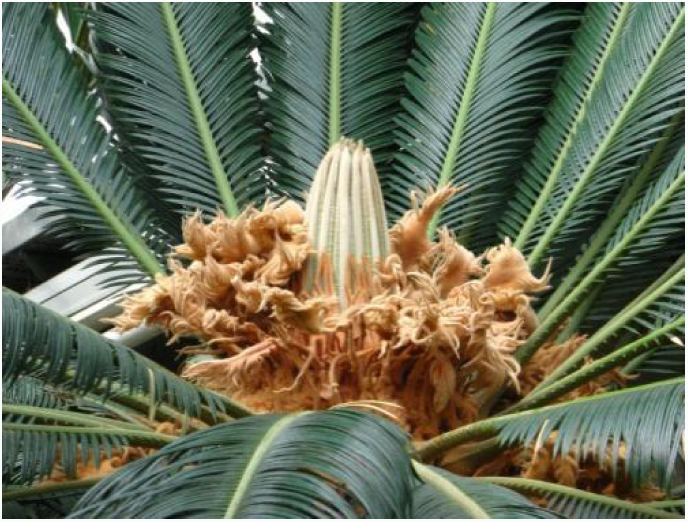
*Cycas revoluta L. Y*oung leaves following female sporophylls of last year.

**Fig. 9B:**
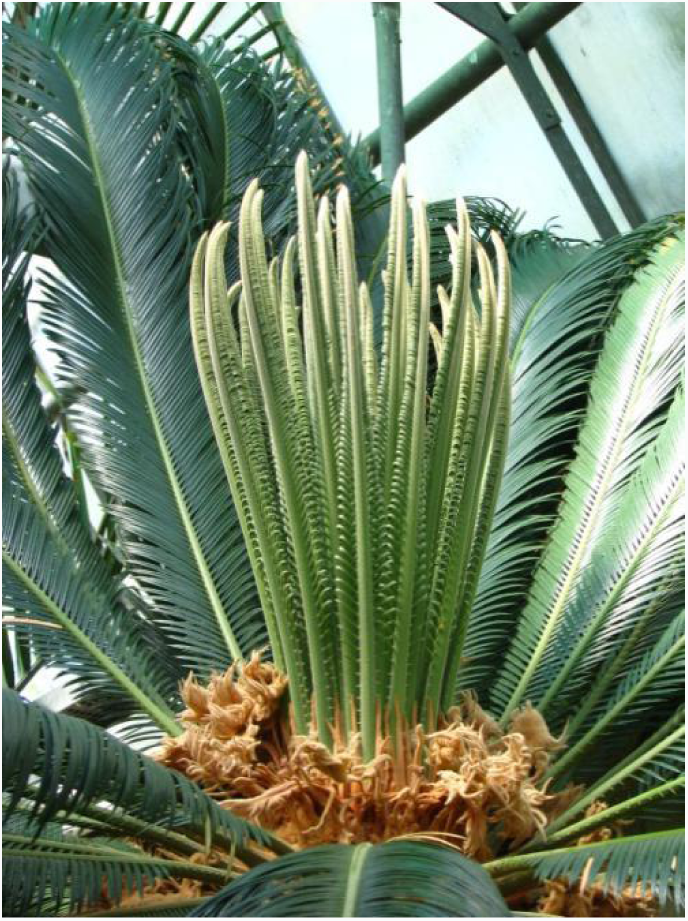
*Cycas revoluta L. Y*oung leaves erecting.

**Fig. 9C:**
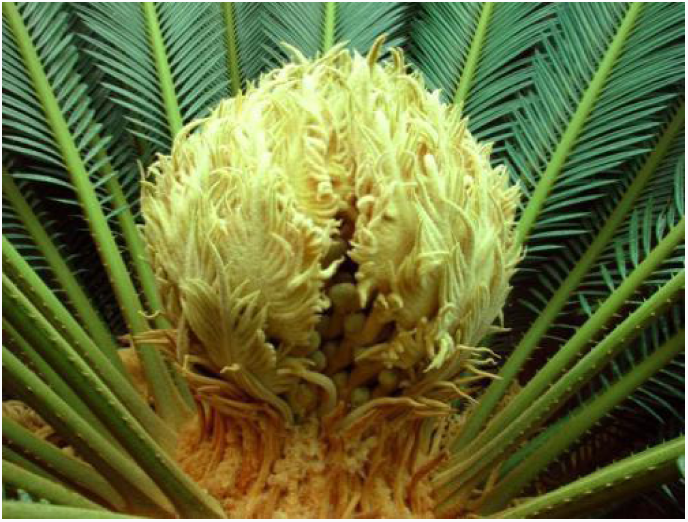
*Cycas revoluta* L. female cone.

**Fig. 10:**
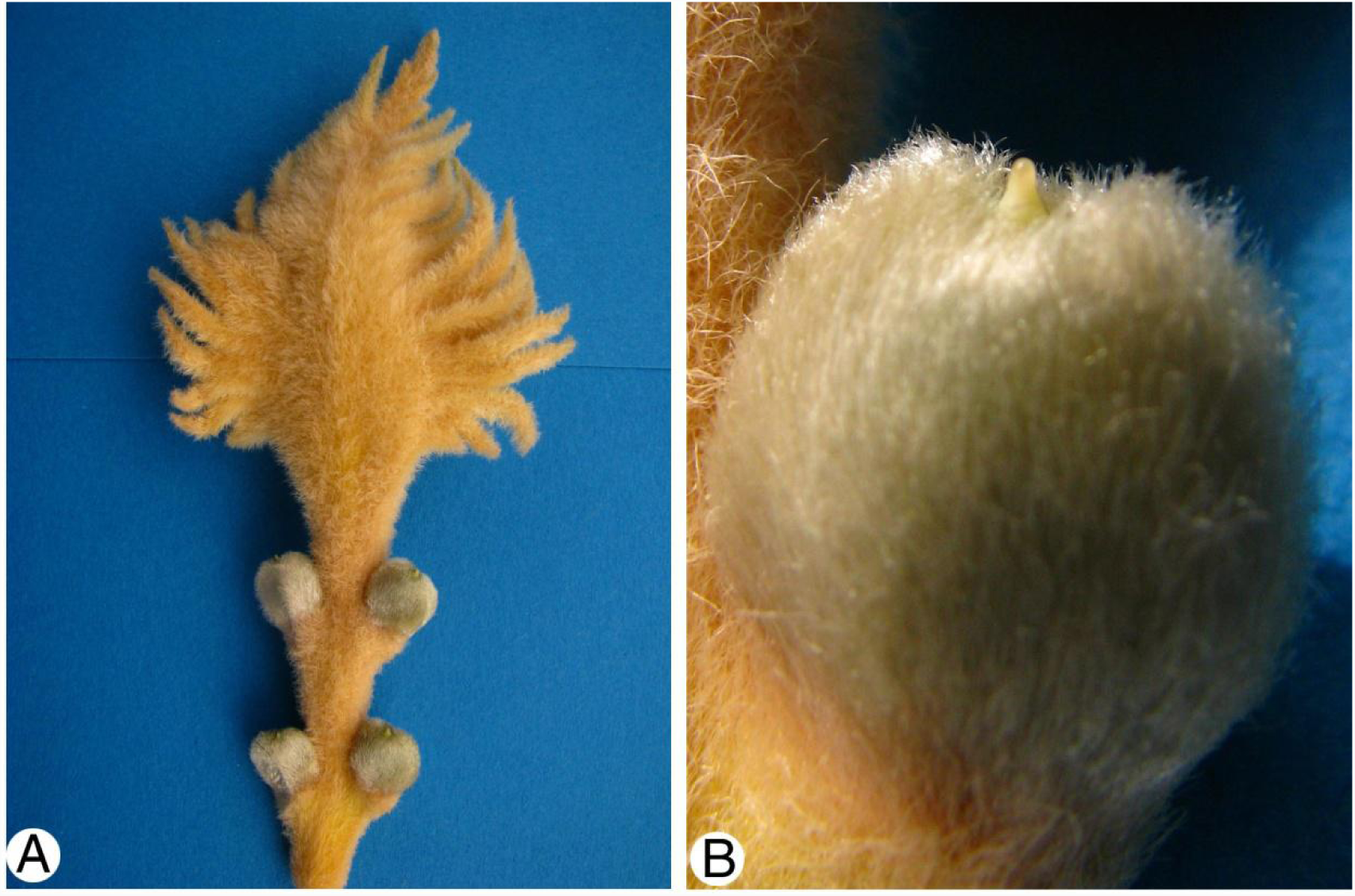
The megasporophyll and ovule of *Cycas revoluta* L. A –The megasporophyll; B – One ovule of the megasporophyll with a protruding receptive surface.

**Fig. 11:**
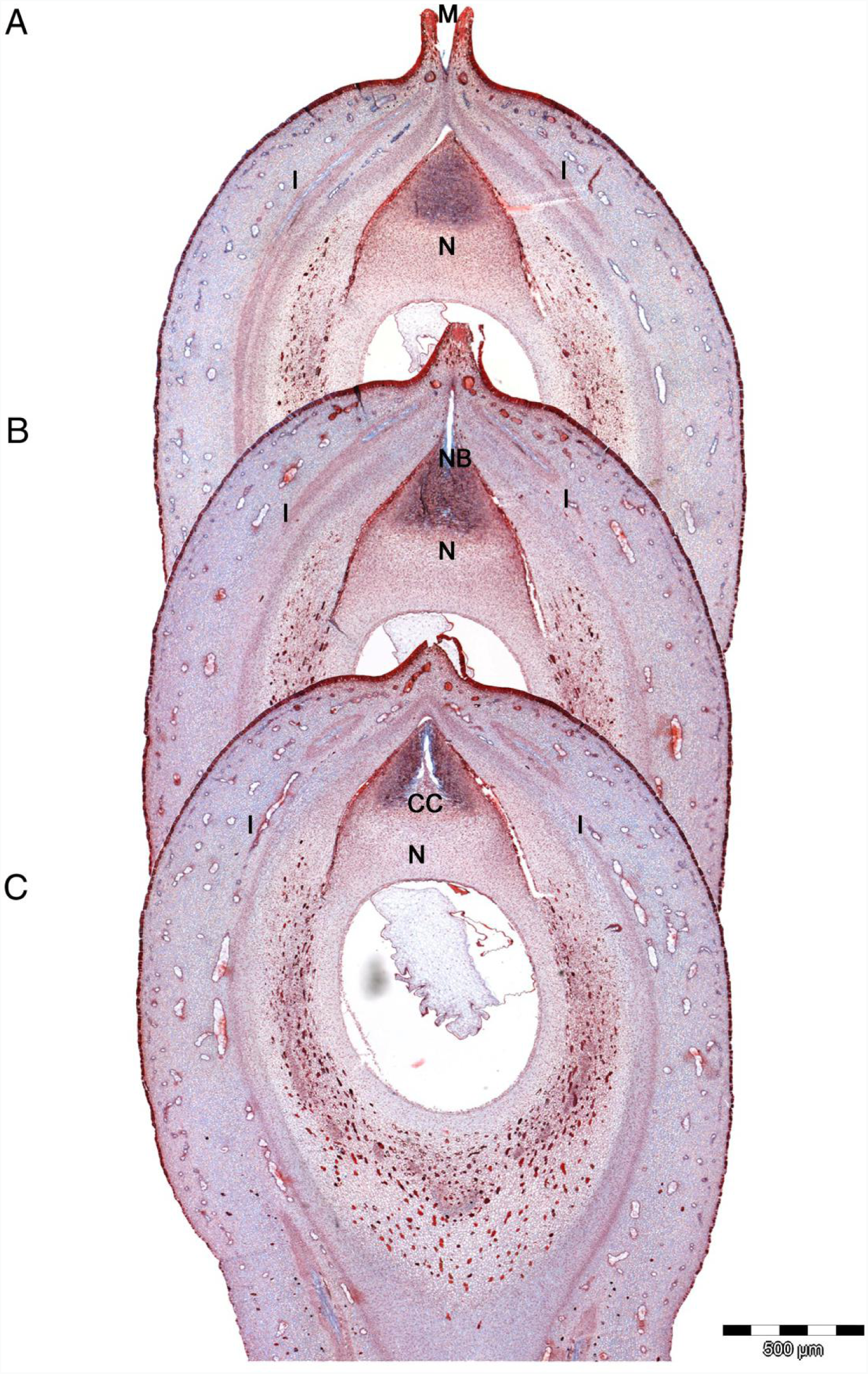
The anatomy of the ovule of *Cycas revolute* L. A, B and C – three longitudinal section of the same ovule; A – micropyle showing; B – nucellus beak showing; C – Central column showing. CC = Central column, I = Integument, M = Micropyle, N = Nucellus, NB = Nucellus beak.

The outer fleshy layer (Fig. 11A, B and C) is called sarcotesta and formed by large, thin-walled undifferentiated cells, filled up with starch. There are many mucilage canals and tannin cells within it. The epidermis is composed of radially elongated cells with thick outer and radial walls, which is very characteristic for the whole genus.

The middle layer (Fig. 11A, B and C) is called sclerotesta which is thickest at the top and near the chalazal end. In the chalazal region it slightly protrudes, accompanying the vascular bundle entering the seed. The cells are thick-walled, and two layers are to be distinguished, an inner one with cells elongated in a longitudinal direction and an outer one with cells directed in a more horizontal plane.

The inner fleshy layer (Fig. 11A, B and C) Is called endotesta which is well developed, strongest at the base and composed of parenchymatous cells, great-cellular, thin-walled, delicate tissue, soon collapsing in the ripe seeds. As in the sarcotesta it contains many tannin cells.

In the micropylar (Fig. 11A) region the sarcotesta forms two thick cushions, with a groove between them, in which the micropyle stands as a little prominent tube. Eichler (1889) says “speaking about the micropyle of Cycas, the seeds I have seen had a micropyle that was always round without any indication of slits. But it is quite possible that other species will show those lobes more distinctly.”

At the free tip of the nucellus is a beak (Fig. 11 B), in which the pollen-chamber is located. This beak has the epidermis of elongated cells, whose outer walls are slightly cuticle covered. The Nucellus tip is solid, and three regions can here be distinguished in the beak. The lower part of the beak is formed by cells which are slightly elongated in the horizontal direction. The cells of the upper part have thick, dark, rich granular content, and in the middle, there is a strand of cells, which are vertically, and in which the formation of the pollen chamber begins. The protruding beak of the nucellus reaches at least into the middle of the micropylar tube. In the center of the protruding beak a longitudinal channel is formed from the tip down to the middle of the nucellar beak. The cells of the beak start to disintegrate when the female cone opens and form a protruding receptive surface.

The formation of the pollination chamber starts with funnel like extension of the beak channel leaving a coniform central column somewhat like the “tent pole” described in Ginkgo (cc in Fig. 11C).

The vascular system consists of three bundles entering the base of the ovule. The median one runs straight on, to the endotesta and divides there into several branches. The two lateral bundles enter the sarcotesta, at the base of which they let off one bundle each, breaking through the sclerotesta and running in the endotesta to the micropyle. The main-bundles continue their way through the sarcotesta to the top.

## Discussions

Reproduction is the most complex part of the life, especially in seed plants. The process of seed plants reproduction starts from the ovule initial. The ovule structure will become different parts of the seed. To understand the function of different structures of a seed, its ontogenies should be investigated. The nucellus of seed plants is one of those questions. The evolutionarypath way of the seed can be just found a clue from the development.The function of the nucellus beak is easy to comprehend aspollination structure of the ovule. However, such tissues have evolved on different organizational levels and in different ways from the outer most tissues.

### Origin of ovule and integument

The widely accepted theory regards the integument as the result of the fusion of sheathing telome (Fig. 12). For this concept usually Andrew is cited, his ideas are however probably based on earlier precursors. In the 19^th^ century was termed “seed buds” in accordance with this. The integument is regarded as a protective structure. However, there have two problems.

**Fig. 12:**
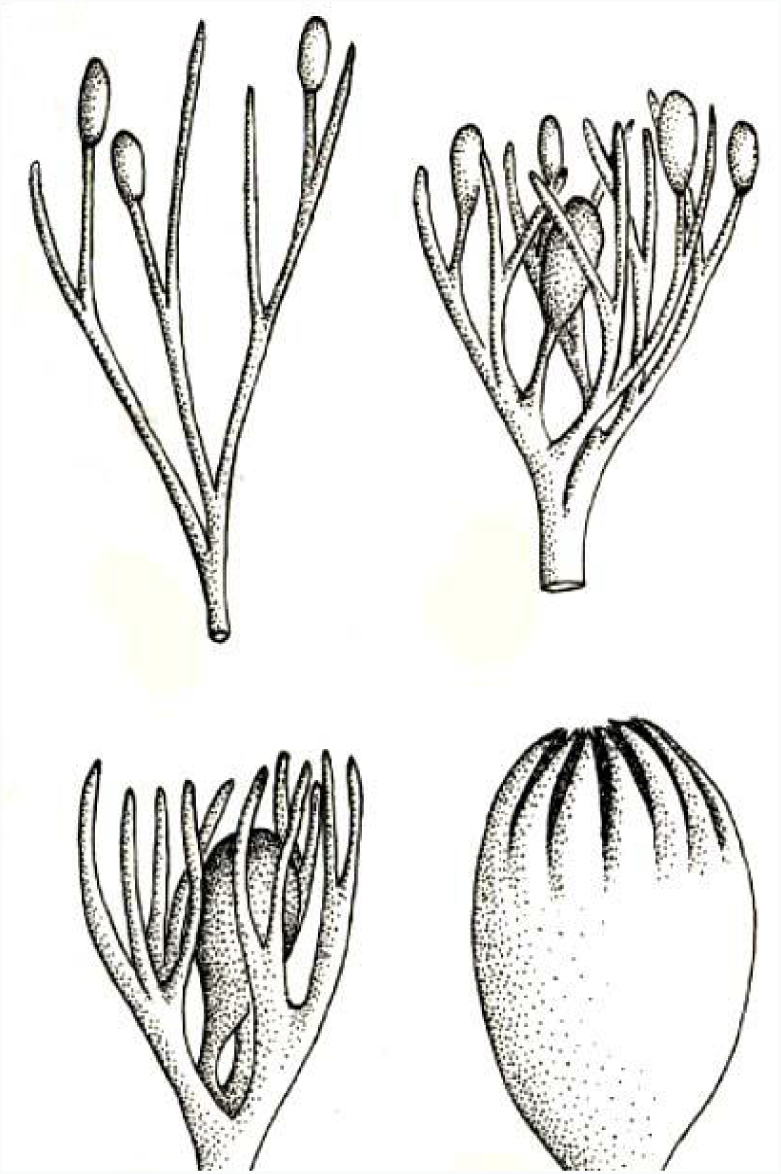
The telome hypothesis after Andrews (1961) from Stewart (1983).

1. At the time of pollination there is still no nutritive tissue to be protected; the function of the micropyle is needed.
2. The development of the integument in gymnosperms does not coincide with the theory in angiosperms.

### Ovules in Lyginopteridatae

The ovule structure of Lyginopteridatae (Fig 13) is a model for the original ovules. However, how do paleobotanist’s distinguish between the two possible interpretations between the ovule of extinct and extant plants? This is a gab for understand the evolution of the seed plants, especially for the evolution of the ovules in seed plants. Moreover, do paleobotanists know the development and function of the „envelopes “. While they claim that the integument of gymnosperms evolved by means of a transgression of function from the salpinx to a cupule like structure. This paper tries the different taxa of basic gymnosperms, *Cycas* and *Zamia*to get some clues of the development and function of the envelopes of ovule and to interpret the evolution of the ovule in seed plants.

**Fig. 13:**
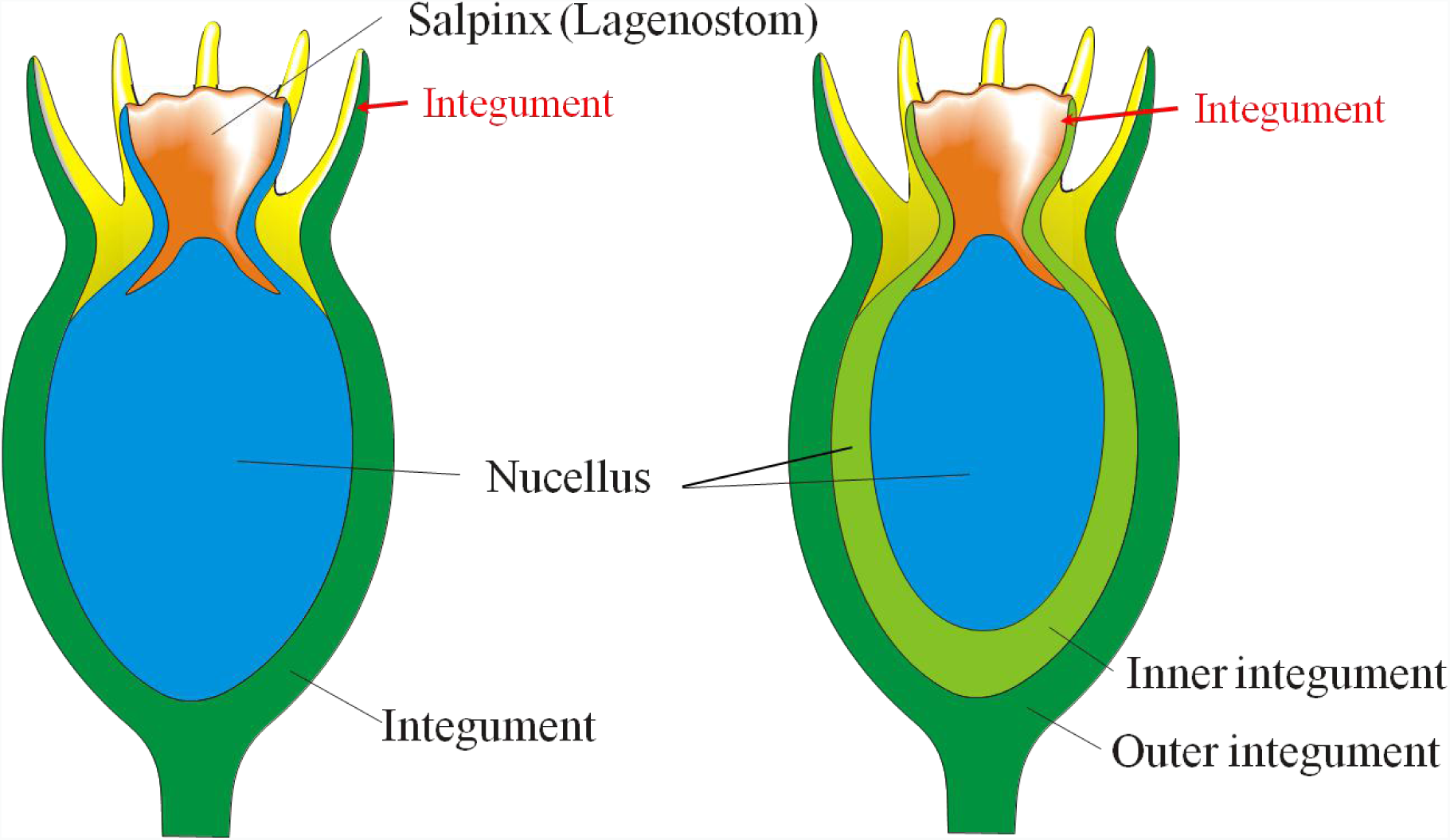
the structure of the Ovules in Lyginopteridatae after Camp & Hubbard (left) modified by Stützel (right).

### The pollination drops

The pollination drop exists in most of the Gymnosperms, except for a few members (*Abies*, *Cedrus*, *Larix*, *Pseudostuga* and *Tsuga*) of Pinaceae, which show a stigmatic micropyle, and a few more (*Araucaria*, *Agathis*, and *Tsuga dumosa*) in which the pollen grains do not land on the micropyle, at the time of pollination (Singh, 1978). Its presence has been mentioned even in the fossil gymnosperms (Andrews 1963). The fluid serves as a receptor of the wind-borne pollen, as also a vehicle for transporting it to the nucellus. It can also find that is a very suitable medium for pollen germination under in vivo condition. (Anhaeusser, 1953; Ziegler, 1959). There are two kinds of pollination drops depending on their formation (Fig. 14 A-D). Chamberlain (1935) and Dogra (1964) and our results (Fig. 14 A and B) showed that for Cycads and Pijl (1953) for Gnetum the pollination drop is formed by breaking down of the cells of nucellusbeak. Ziegler (1959) concluded that the osmotically active substances released by the degenerating cells (which may contain druses) absorb water from the nucellus or the atmosphere, fill the micropylar canal and is seen as a drop at the micropyle. The other kind of pollination drop in those plants (e.g. cupressads) in which the nucellar cells do not degenerate before pollination. Doyle and O’Leary (1953) and McWilliam (1958) working on Pinus, and Ziegler (1959) working on *Taxus*, stated that the drop is secreted by the cells forming the apex of the nucellus. This is like the results which I found in *Cupressus* (Fig. 14 C and D). What the different composition and the connection between those two different pollination drops are not so clear until now. How is the function and evolution of the pollination to merit further investigation?

**Fig. 14:**
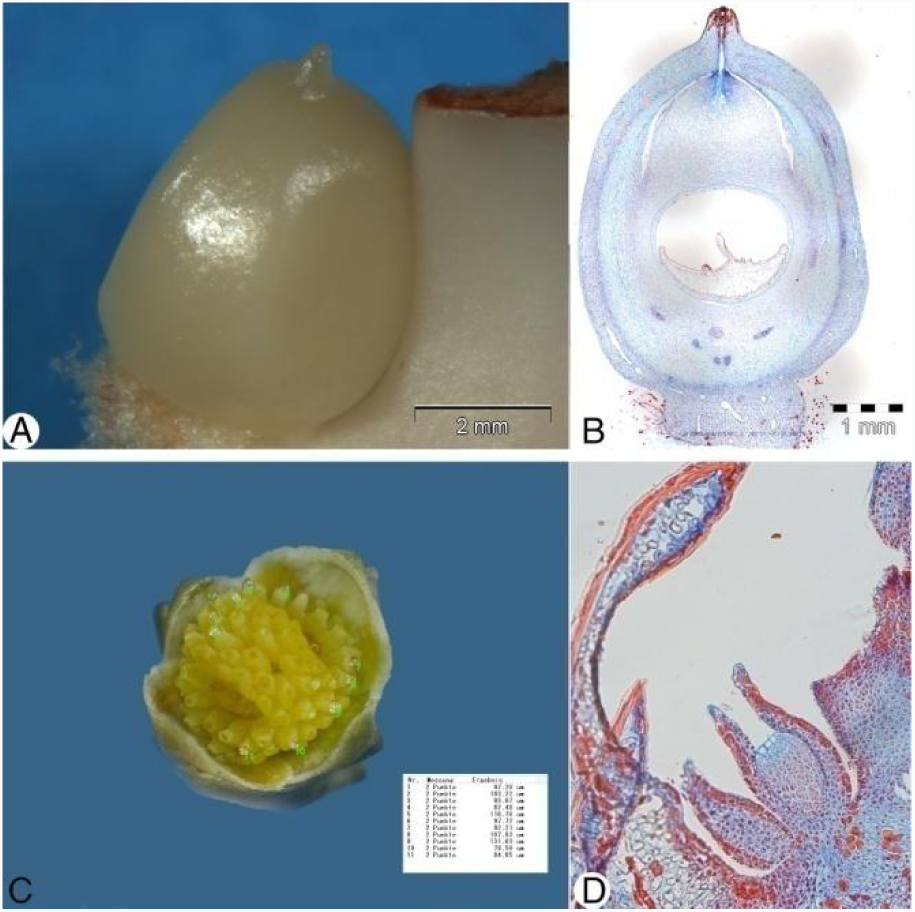
Pollination drop of *Zamia amblyphyllidia* and *Cupressus arizonica*. A.-the pollination drops of *Z. amblyphyllidia*; B. -the longitude section of Zamia ovule in A; C. - the pollination drops of *Cupressus arizonica*; D. - the longitude section of the ovule of *Cupressus arizonica*.

### The pollination of *Cycads*

Chamberlain (1919) considered that *Cycas* is wind-pollinated plant. However, there are also reports of *Encephalartos* and *Macrozamia* are insect pollination plants. Oberprieler (1995) recorded the life cycle of 63 weevil specise, belonging to 12 genera, associated with cycads and even regarded that ants can be the pollinator of *Cycas*.

There are two processes of the pollination of the cycads. First, the pollen needs to transport from the microsporophyll to the microsporophyll, whatever by wind or insect. Then there also need some of the medium to transport the pollen from the microsporophyll to the pollination drop. Norstog (1990) considered that the insect probably is the pollinator of the *Dioon* and *Zamia*, and *Cycas* need the fluid to bring the pollen from megasporophylls to the ovule, after his simulated experiment of the wind pollination of *Dioon*, *Zamia* and *Cycas*.

In our results, before pollination forming, the cell of the nucellar beak of *Zamia* will start deliquescing (Fig. 7A). It will be produced like a pollination drop hanging on the micropyle. It will be happened from the top of the nucellar beak to the base of the beak. There will have just a circle cell wall around the micropylar tube (Fig. 7A, C and E). This moment, if there is any pollen into the strobilus from the opening. This liquid tissue will stick the pollen. Then the nucellus will absorb the liquid into the micropylar tube (Fig. 7C, D). Later the pollens will site on the pollen chamber. At this time the multinucleated Macro prothallium developed into a multicellular tissue (Fig. 7D).

On the other hand, artificial pollination under greenhouse conditions are performed in *Zamia* by using water with dispersed pollen through the strobilus. Freely exposed pollination droplets would be mashed away applying this method. So, the pollination drop of Cycads is different from the normal pollination drop of conifers which is secreted by the cells forming the apex of the nucellus. The pollination drop of cycads is formed by breaking down of the cells of nucellar beak. This process is the same as fossil gymnosperms (Andrews, 1963).

. In living gymnospermous seed plants, the pollen is caught by a pollination drop exuded from micropyle, but this was impossible before fusion of the integument lobes. Instead the pollination drop was exuded by an extension of the nucellus, called the lagenostome or salpinx, in which the epidermis separated from the inner tissue to form a central column surrounded by a pollen chamber (hydrasperman reproduction: Rothwell & Scheckler 1988).

### The ovule and nucellus comparison between *Cycas* and *Zamia*

The ovule of Cycas is bilateral symmetrical and covered near the micropyle with hairs, whereas young ovules are wholly clothed by a dense hair covering except the micropyle. The micropyle points to the upside of the strobilus. There is a special structure on the pollination chamber of *Cycas*. Similar structure is called tend-pole in *Ginkgo biloba* by and central column in *Lagenostoma* (Long, 1960).

The ovule of *Zamia* is radial symmetrical and bare without hair covering. The micropyle points to the axis of the strobilus and slightly towards one another. The ovules are clearly not marginal, but each is situated slightly within the margin of a small area apparently corresponding to one of the areas of the lower surface of a female sporophyll.

According to Stopes (1904, 1905), these two sarcotesta bundles are the remnants of the original radio-symmetry and many-bundles condition of the outer flesh. The female gametophyte is very large.

### The sperm and fertilization

We got the special structure on the top of one archegonium (Fig. 4C, D). This structure looks like a sperm sit on the top of the neck cells. There is a structure outside this cell looking like the branch of the flagella. However, normally it is considered that this is a structure of the neck cells which is broken with some fragments on the out circle of the cell. The fertilization of the *Cycas* and *Gingko* is not so clear today. Even the sperm will be totally in to the egg or not is still a question. After pollination, the development of pollen from the beginning to two functional flagella sperms will take four or five months. Nevertheless, the fertilization will be finished on several hours. So, it is difficult to hit the point of the fertilization.

### Conclusion

The nucellar beak found in *Zamia* is a structure that has not been recorded previously. It protrudes from the micropyle at pollination and may be the primary acceptor for pollen. There are striking similarities to the Lagenostoma or salpinx in Lyginopteridatae. There may be an evolutionary way to interpret the pollination drop existing in the Lyginopteridatae. Probably the nucellar beak of Cycads, even Ginkgoales have the same function with the Lagenostoma or salpinx of the Lyginopteridatea. Unfortunately, pollen and transport inside the pollination chambers have not been observed. Further analysis of this unusual structure seems to be very important.

## Acknowledgment

The Botanical Garden of the Ruhr University Bochum has generously allowed me to collect the material for the current and several other studies. The first author is especially grateful to Prof. Thomas Stützel who give me some much support of this project, Sabine Adler for her efforts in linguistically improving the manuscript and Patrick Knopf of helping with illumination. Special acknowledgment is due to Dr. Christian Schulz who read the manuscript and gave us very useful advices. This work was financially supported by he Research Funds for the Doctoral Program of Shaanxi Province (A279021830) to X. Z. and the Fundamental Research Funds for the Central Universities (Z109021614) and the Research Fund for the Doctoral Program of Higher Education of China (Z109021501) and the Chinese Scholarship Council.

## References

Albert G. Long II.—On the Structure of Calymmatotheca kidstoni Calder (emended) and Genomosperma latens gen. et sp. nov. from the Calciferous Sandstone Series of Berwickshire

Albert G. Long XII.—On the Structure of Samaropsis scotica Calder (emended) and Eurystoma angulare gen. et sp. nov., Petrified Seeds from the Calciferous Sandstone Series of Berwickshire

Albert G. Long IX.—Stamnostoma huttonense gen. et sp. nov.—a Pteridosperm seed and cupule from the Calciferous Sandstone Series of Berwickshire

Andrews, H. N. 1961 Studies in paleobotany. New York.

Andrews, H. N. 1963 Early seed plants. Science 142: 925–931.

Anhaeusser, H. (1953): Keimung und Schlauchwachstum des Gymnospermenpollens unter besonderer Berueksichtigung des Wuchsstoffproblems. Beitr. Biol. Pfl. 29: 297–338.

Benson, M. 1904. Telangium scottii, a new species of Telangium (Calymmatotheca) showing structure. Annals of Botany 18: 161–176.

Camp, W., & Hubbard, M. (1963). On the Origins of the Ovule and Cupule in Lyginopterid Pteridosperms. American Journal of Botany, 50(3), 235–243. Retrieved from http://0-www.jstor.org.explore.searchmobius.org/stable/2440016

Chamberlain, C. J. 1919. The living Cycads. The University of Chicago Press, Chicago, Illinois.

Chamberlain, C. J. (1935): (reprint 1966). Gymnosperms: structure and evolution. –Chicago Univ. Press, Chicago.

Dogra, P. D. (1964): Pollination mechanisms in gymnosperms, - In P. K. K. Nair (Ed.): Recent Advances in Palynology. 142–175. National Botanic Gardens, Lucknow.

Doyle, J. and O’Leary, M. (1935): Pollination in Pinus. – Sci. Proc. Roy. Dublin Soc. 21: 181–190.

Eames, A. J. (1961). Morphology of angiosperms. New York.

Gensel, P. G. and Andrews, H. N. 1984 Plant life in the Devonian. Praegler, New York.

Haan, H. R. M. De 1920 Contribution to the knowledge of the morphological value and phylogeny of the ovule and its integuments. Rec. Trav, bot, néerl. 17 219–324.

Herr, J. M., Jr. 1995 the origin of the ovule. American Journal of Botany 82(4) 547–564.

Jones, D. L. 1993 Cycads of the world. Reed Books, Chatswood.

Long, A. G. 1966 some lower carboniferous fructications from Berwickshire, and carpels. Transactions of the Royal Scientific Proceedings of the Royal Dublin Society 23:35–54.

NORSTOGK. 1990. Studies of cycad reproduction at Fairchild Tropical Garden. Mem. New York Bot. Gard. 57:63–81

Matte, M. 1904 L’appareil libéro-ligneux des Cycadacées. Caen.

McWilliam, J. R (1958): The role of the micropyle in the pollination in Pinus. - Bot. Gae. 120: 109–117.

Meeuse ADJ, Bouman F. The inner integument – its probable origin and homology, Acta Botanica Neerlandica, 1974, vol.23 (pg.237–249)

Oberprieler RG 1995 The weevils (Coleoptera: Curculionoidea) associated with cycads. 1. Classification, relationships, and biology. Pages 295–365 in P. Vorster, ed. Proceedings of the third international conference on cycad biology. Cycad Society of South Africa, Stellenbosch.

Oliver, F. W. and Scott, D. H., 1904 On the structure of the Paleozoic seed *Lagenostoma lomaxi*, with a statement of the evidence upon which it is referred to *Lyginodendron*. Philosophical transactions of the Royal Society, London 197B: 193–247.

Pijl, V. van der (1953): On the flower biology of some plants from Java. – Ann. Bogor. 1: 77–99.

Rothwell, G.W. & Scheckler, S.E. (1988) Biology of ancestral gymnosperms. In: Origin and Evolution of Gymnosperms (ed. C.B. Beck). pp. 85–134. Columbia University Press, New York.

Singh, Hardev 1978 Embryology of Gymnosperms Encyclopedia of Plant Anatomy. Gebrüder Botntraeger, Berlin, Stuttgart.

Singh, H 1978 Embryology of Gymnosperms -Gebrueder Borntraeger, Berlin, Stuttgart

Smith, D. L. 1964 The evolution of the ovule Biological Review 39: 137–159.

Stevenson, D. W. 1987. Comments on character distribution, taxonomy, and nomenclature with respect to the genus Zamia L. in the West Indies and Mexico. Encephalartos 9: 3

Stopes, M. C. 1904 Beiträge zur kenntnis der fortplanzungsorgane der Cycadeen. Flora, 93, 435–482.

Stopes, M. C. 1905 On the double nature of the Cycadean integument. Annals of Botany Vol. XIX. No. LXXVI.

Taylor, T. and Taylor, E. 1993 The biology and evolution of fossil plants, Prentice Hall, Englewood Cliffs, NJ.

Taylor, T., Taylor, E. and Krings, M., 2009 Paleobotany-the biology and evolution of fossil plants second edition Elsevier.

Walton, J. 1953 the evolution of the ovule in the pteridosperms. Advancement of Science 38: 223–230.

William H. Lang, 1900 studies in the development and morphology of Cycadean Sporangia: II the ovule of *Stangeriaparadoxa*, Annals of Botany Vol 14

Worsdell, W. C. 1904. The structure and morphology of the ovule: an historical sketch. Annals of Botany 18: 57–86.

Ziegler, H. (1959): Ueber die Zusammensetzung des “Bestaeubungstropfens” und dem Mechanismus seiner Sekretion. –Planta (Berlin) 52:587–599.

Zimmermann, W. 1938 Die Telomtheorie. Biologe: Monatsschrift zur Wahrung der Belange der Deutschen Biologen 7: 385–391.

Zimmermann, W. 1952 Main results of the telome theory. Palaeototanist, 1, 456–470.

